# The ER transmembrane complex (EMC) can functionally replace the Oxa1 insertase in mitochondria

**DOI:** 10.1101/2021.08.02.454725

**Authors:** Büsra Güngör, Tamara Flohr, Sriram G. Garg, Johannes M. Herrmann

## Abstract

Two multisubunit protein complexes for membrane protein insertion were recently identified in the endoplasmic reticulum (ER): The guided entry of tail anchor proteins (GET) complex and ER membrane complex (EMC). The structures of both of their hydrophobic core subunits, that are required for the insertion reaction, revealed an overall similarity to the YidC/Oxa1/Alb3 family members found in bacteria, mitochondria and chloroplasts. This suggests that these membrane insertion machineries all share a common ancestry. To test whether these ER proteins can functionally replace Oxa1 in yeast mitochondria, we generated strains that express mitochondria-targeted Get2-Get1 and Emc6-Emc3 fusion proteins in Oxa1 deletion mutants. Interestingly, the Emc6-Emc3 fusion was able to complement an *Δoxa1* mutant and restored its respiratory competence. The Emc6-Emc3 fusion promoted the insertion of the mitochondrially encoded protein Cox2 as well as of nuclear encoded inner membrane proteins though was not able to facilitate the assembly of the Atp9 ring. Our observations indicate that protein insertion into the ER is functionally conserved to the insertion mechanism in bacteria and mitochondria and adheres to similar topological principles.

## Introduction

Membranes of bacteria and eukaryotic cells contain different protein translocases. These pore-like structures transport unfolded polypeptides across membranes and, in case of membrane proteins, laterally integrate them into the lipid bilayer [1]. Examples are the SecY/Sec61 complexes of the bacterial inner membrane and the endoplasmic reticulum (ER)[2, 3], the beta barrel-structured outer membrane translocases of bacteria, mitochondria and chloroplasts [4, 5], and the translocases of the mitochondrial inner membrane (TIM23 and TIM22 complexes) [6, 7]. These translocases belong to distinct non-related protein families and developed independently during evolution.

Protein translocation can also be mediated by a second group of translocation machineries which do not form defined pores but rather facilitate protein translocation by local distortion and compression of lipid bilayers [8]. Such a mechanism was recently proposed for the ER-associated degradation (ERAD) pathway machinery [9].

Locally distorted and compressed lipid bilayers are also used by insertases, which integrate hydrophobic proteins into membranes. Substrates of these insertases include membrane proteins that lack large hydrophilic regions on the trans-side of the membrane (such as in the case of tail-anchored proteins) or multispanning membrane proteins whose more complex topogenesis relies on the cooperation of insertases with a canonical protein translocase.

The mitochondrial protein Oxa1 was discovered in the early 90s as the first representative of these insertases [10, 11] and served as the founding member of the YidC/Oxa1/Alb3 family. These closely related and functionally exchangeable proteins [12-15] mediate membrane insertion of proteins into the inner membranes of bacteria and mitochondria as well as in the thylakoid membrane of chloroplasts [16-23]. Several YidC structures were published recently which suggest that these monomeric proteins serve as enzymes that accelerate the spontaneous (though often inefficient, slow and non-productive) partitioning of hydrophobic segments into the lipid bilayer [24-27].

Two recently identified protein complexes serve as insertases in the ER membrane: The Get1-Get2 (in vertebrates WRB-CAML) complex which facilitates the insertion of tail-anchored proteins [28] and the ER membrane complex (EMC), a multimeric insertase consisting of eight (yeast) or nine (animals) subunits which acts independently of as well as in conjunction with the Sec61 translocon in the topogenesis of multispanning ER proteins [29-33]. The structures of both complexes were recently solved [34-36], revealing a striking similarity of the reaction centers formed by Get1-Get2 and Emc6-Emc3 with the architecture of YidC/Oxa1/Alb3 proteins despite very limited sequence similarity. On the basis of their structural architecture, it was proposed that all these insertases are members of one related group of proteins, which was named the Oxa1 superfamily [37].

In this study, we report that mitochondrial Oxa1 protein can be functionally replaced by the core components of the EMC complex, at least in respect to its role in the membrane insertion of proteins. Unlike Oxa1, EMC is however unable to facilitate the assembly of the F_o_ sector of the ATPase, presumably because it is not recognized by mitochondrion-specific assembly factors [38, 39]. Our study shows that the EMC complex of the ER and the Oxa1 insertase of mitochondria fulfill analogous molecular functions, consistent with their proposed structural similarity.

## Results

### The core components of the various membrane insertases display similar topology despite limited sequence identity

The YidC, Alb3 and Oxa1 insertases of bacteria, chloroplasts and mitochondria are characterized by five conserved transmembrane domains (Fig. 1A). The DUF106 protein family of archaea was proposed to be a distant relative, although it only has three transmembrane domains which show similarity to the transmembrane domains 1, 2 and 5 of Oxa1. It was suggested that the DUF106 family gave rise to Emc3 and Get1 on the basis of very similar overall structural organization [34, 35].

**Fig 1.**
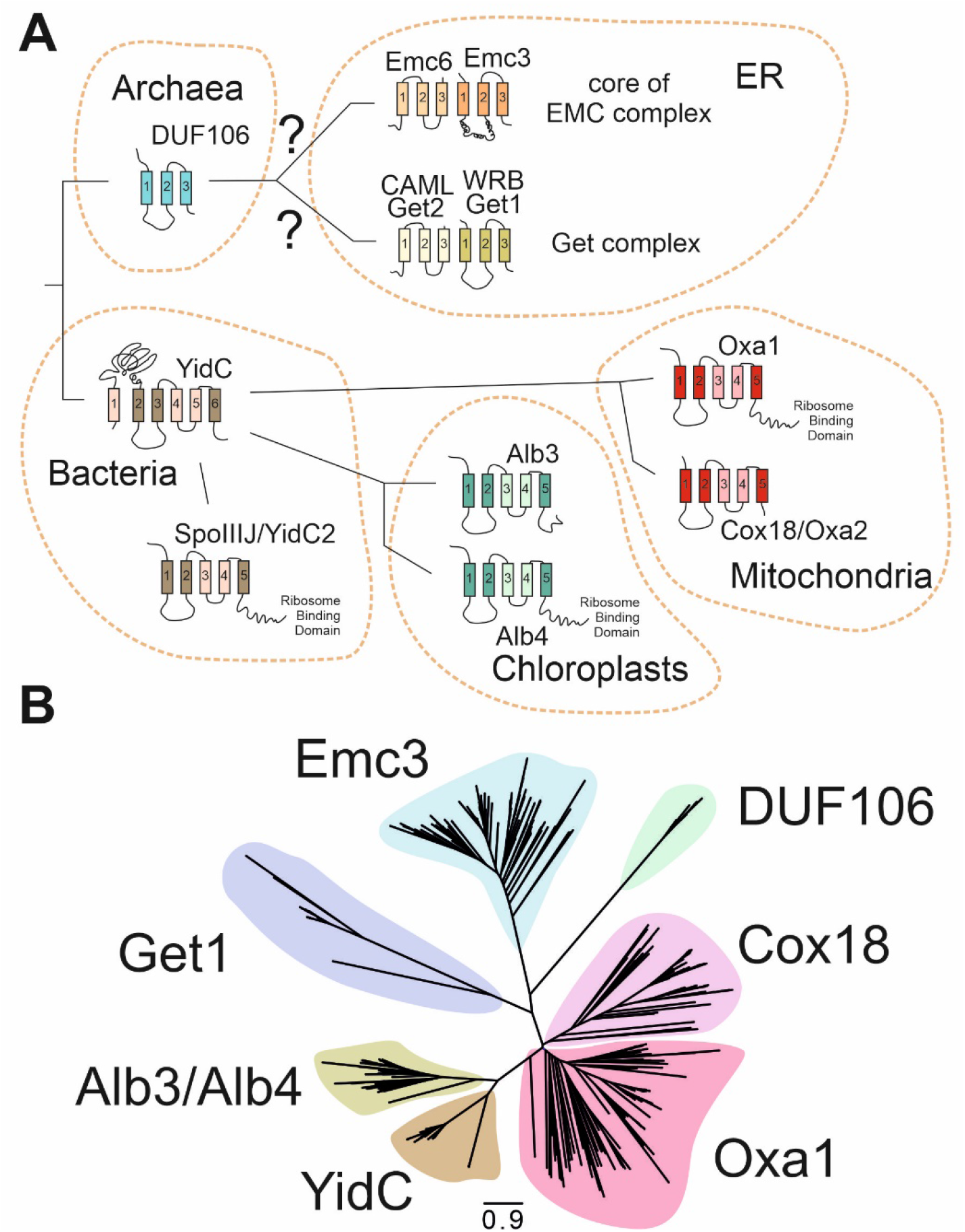
The members of the Oxa1 superfamily are structurally similar, however, their phylogenetic relationship is unclear. (**A**) All shown proteins serve as membrane insertases. Whereas the representatives of bacteria, mitochondria and chloroplasts are monomeric proteins with five or six transmembrane domains, the insertases of the ER form oligomeric complexes of two (Get1/Get2) or multiple (EMC) subunits. Recent studies on the structure of the GET and EMC complexes proposed that Emc3 and Get1 are structurally related to YidC, Oxa1 and Alb3. The transmembrane domains that share structural similarity are indicated in darker color. (**B**) Blast searches identified 460 homologs of DUF106, Emc3, Get1, Alb3/Alb4, Oxa1 and YidC proteins from different species (see Fig. S1 and Supplementary Data Files 1-3). This tree supports the relatedness of these protein groups and is consistence with the hypothesis that Emc3 is related to a DUF106-like ancestor protein. While YidC/Oxa1/Alb3 proteins form a closely related group, the branches to Get1, Emc3 and DUF106 are much longer and, thus, their sequences are likely more deviated.

To assess a potential phylogenetic relationship among these proteins, we screened for potential related proteins of Oxa1, Alb3, YidC, DUF106, Emc3 and Get1 and identified 460 unique homologs across eukaryotes and prokaryotes (see supplemental data set 1-3). Phylogenetic trees were calculated based on trimmed alignments (Fig. 1B). These trees supported the common origin of Oxa1, Alb3 and YidC very well and also indicated good bootstrap support for their relationship with members of the DUF106, Get1 and Emc3 families (Fig S1, supplemental data set 1-3). Even though the similarity of individual proteins is low, the inclusion of the large number of sequences allowed the construction of a well-supported tree that supports the relatedness of these different groups of insertases. Even if analogy based on convergent evolution cannot be formally excluded, homology based on common ancestry appears more likely.

### A mitochondria-targeted EMC core restores respiration of *Δoxa1* cells

The recently solved structures of the EMC and GET complexes [34-36] suggested that their core centers, consisted of Emc6/Emc3 and Get2/Get1 respectively, resembling the structural organization of YidC. This inspired us to clone the respective regions of Emc6/Emc3 and Get2/Get1 into fusion proteins which also contained the matrix-targeting sequence (MTS) of Oxa1 to ensure mitochondrial import of these proteins, the C-terminal ribosome binding domain of Oxa1 necessary for its interaction with the mitochondrial translation machinery [40-43], as well as a short Oxa1-derived linker to confer insertion into the inner membrane (Fig. 2A). However, all transmembrane regions of these fusion proteins were derived from ER proteins.

**Fig 2.**
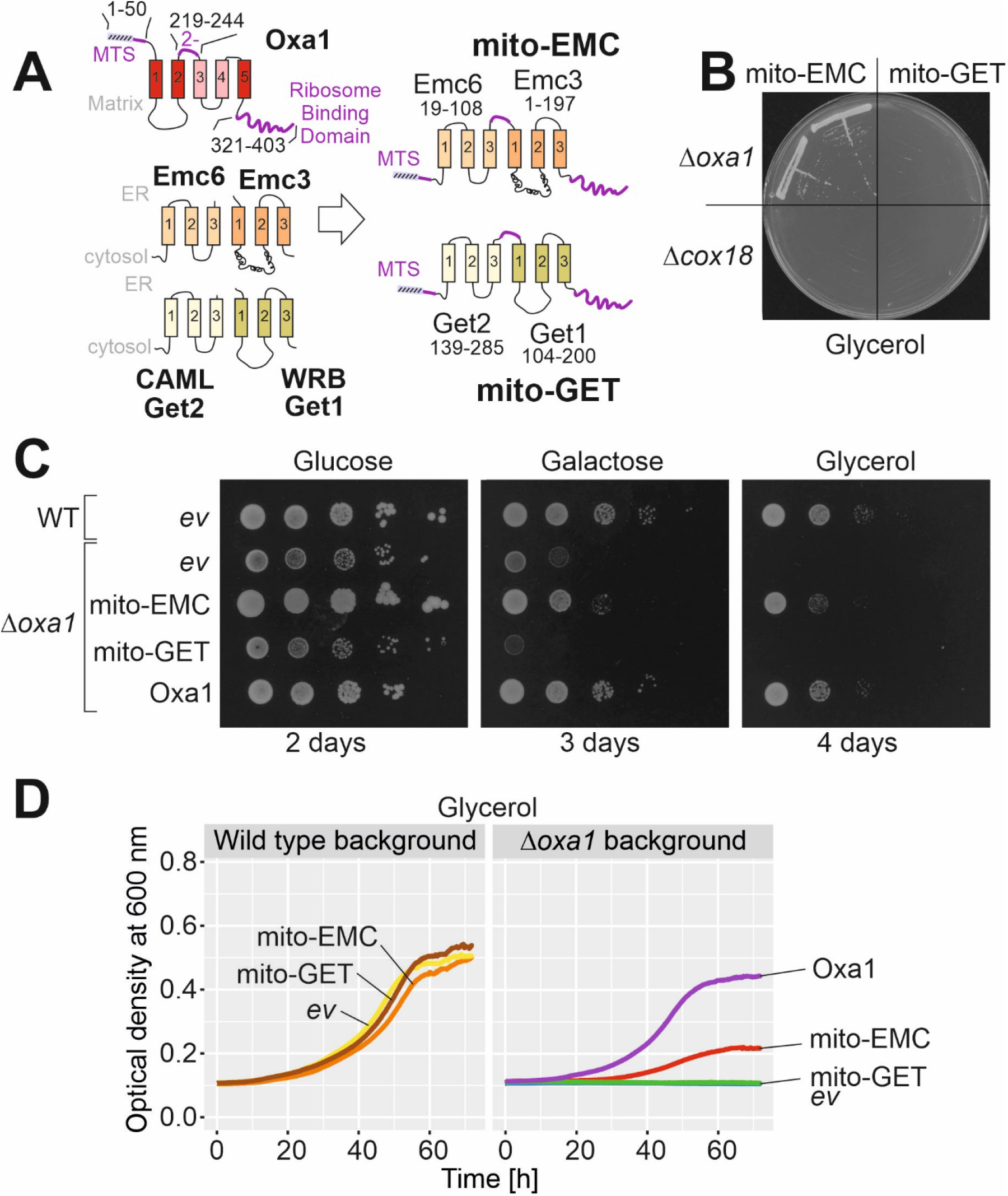
Expression of a mitochondria-targeted Emc6-Emc3 fusion protein (mito-EMC) suppresses the growth defect of a *Δoxa1* mutant. (**A**) Schematic representation of the fusion proteins used in this study. The segments shown in purple are derived from Oxa1 in order to ensure import and insertion of the proteins into the inner membrane. (**B**) Simple growth test on glycerol plates on which cells can only grow if they generate energy from respiration. (**C**) Cells were grown in galactose medium, from which tenfold serial dilutions were dropped onto plates with the respective carbon sources. An empty vector (*ev*) control is shown for comparison. (**D**) Growth curves of *Δoxa1* cells with the indicated expression plasmids. Shown are mean values of three technical replicates (n=3).

These fusion proteins, which we named mito-EMC and mito-GET, were expressed in *Δoxa1* and *Δcox18* cells, thus in mutants lacking Oxa1 or its paralog Cox18. Surprisingly, the expression of mito-EMC partially suppressed the growth defect of *Δoxa1* mutants even upon growth on the non-fermentable carbon source glycerol (Fig. 2B-D) but not that of *Δcox18* cells (Fig. 2B, S2A). We did not observe any suppression by expression of the mito-GET protein suggesting this protein might not be properly integrated into the inner membrane or generally is unable to fully replace mitochondrial insertases in function.

### Mito-EMC can facilitate the insertion of nuclear encoded inner membrane proteins

Oxa1 mediates the insertion of nuclear encoded inner membrane proteins that use the so-called conservative insertion pathway (Fig. 3A). In order to test whether mito-EMC can take over this function of Oxa1, we isolated mitochondria from *Δoxa1* cells that expressed mito-EMC or Oxa1 for control. Radiolabeled precursor proteins of different model substrates were incubated with these mitochondria to allow their import and intramitochondrial sorting. Then, mitochondria were re-isolated and treated with proteinase K to remove non-imported material or converted to mitoplasts by hypoosmotic rupturing of the outer membrane (swelling) and treated with proteinase K. Radiolabeled proteins that were integrated into the inner membrane became protease-accessible giving rise to characteristic fragments (Fig. 3B, 3C, S2B, S2C; white arrowheads). From the ratio of these fragments to the protease-inaccessible species (black arrowheads), the insertion efficiency of these membrane proteins can be estimated (Fig. 3D). These *in vitro* import experiments showed that mito-EMC (and Oxa1) facilitated membrane insertion of the model proteins Oxa1 [44] and Su9(1-112)-DHFR [45]. Previous studies have shown that the Oxa1-mediated insertion efficiency is dependent on the negative charge in the inserted region and that positively charged regions are not exported by Oxa1 [46]. The same charge dependence was also found for mito-EMC-mediated insertion, suggesting that both proteins facilitate membrane insertion by a comparable mechanism that drives the negatively charged sequence to the positively charged side of the inner membrane [27, 47].

**Fig 3.**
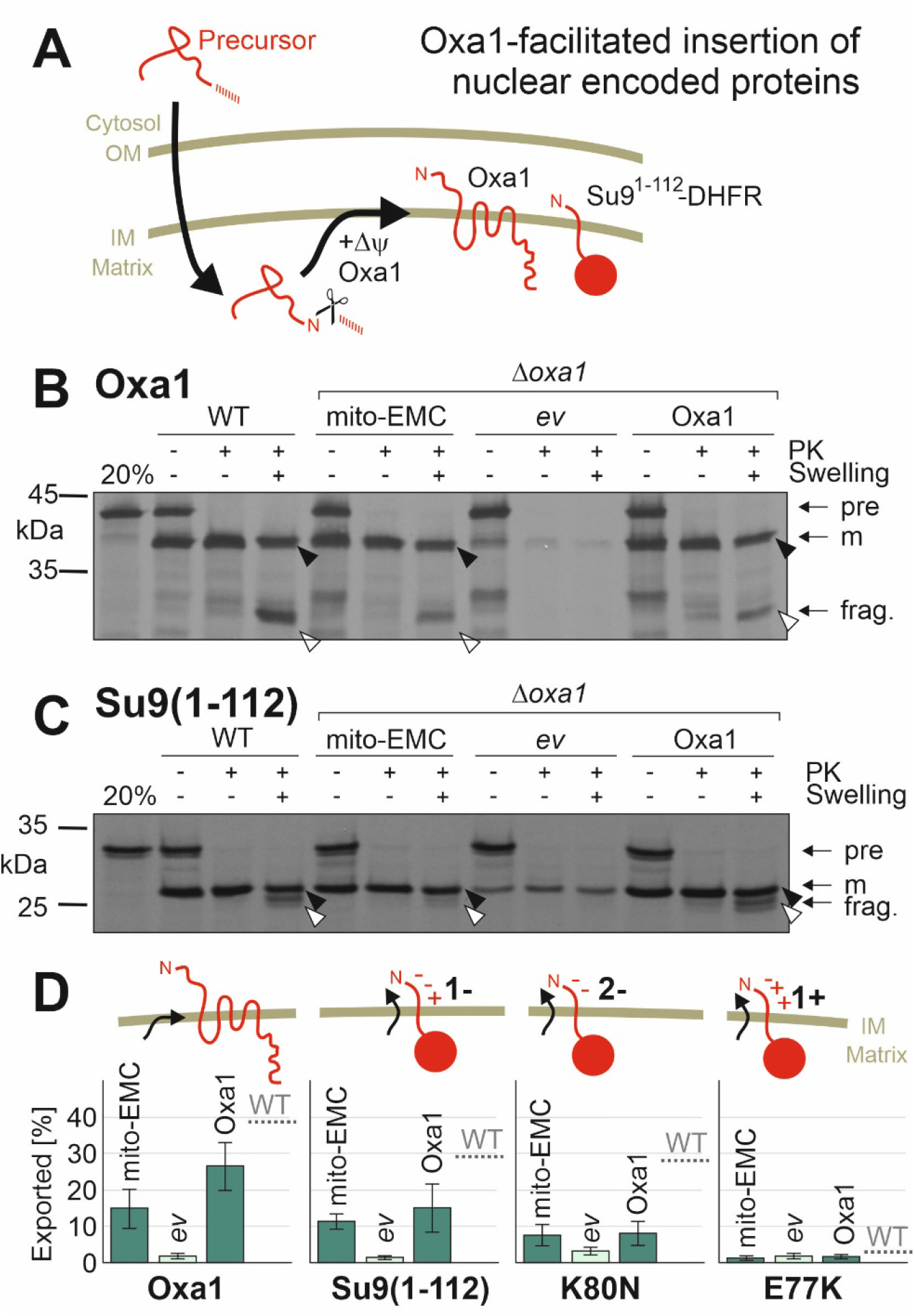
Protein insertion by mito-EMC, like that by Oxa1, depends on negative charges in the transferred sequence. (**A**) Schematic representation of the import and Oxa1-mediated membrane insertion of nuclear encoded proteins. The scissors indicate the proteolytic removal of the N-terminal mitochondrial targeting sequence by the matrix processing peptidase. Protein insertion depends on Oxa1 and the membrane potential (Δψ). (**B, C**) Radiolabeled Oxa1 and Su9(1-112)-DHFR were incubated with mitochondria isolated from the indicated mutants for 30 min at 25°C. Samples were divided into three fractions and treated with or without proteinase K (PK) under isosmotic or hypoosmotic (Swelling) conditions. Upon swelling, protease treatment generates a fragment (frag., white arrowhead) from the membrane-embedded protein, but not from the translocation intermediate that still resides in the matrix (black arrowhead). (**D**) Insertion efficiencies were quantified from three biological replicates and quantified. Mean values and standard deviations are shown.

### Mito-EMC can facilitate the membrane insertion of mitochondrial translation products

Next, we assessed the insertion of mitochondrial translation products. First, we monitored mitochondrial protein synthesis in whole cells after inhibition of cytosolic translation by cycloheximide (Fig. 4A). This showed that *Δoxa1* cells that contained Oxa1 or mito-EMC were able to synthesize proteins, whereas no translation products were observed in cells expressing mito-GET, indicative for the loss of mitochondrial DNA in these cells. The increased depletion of mitochondrial DNA was reported before in Oxa1-deficient cells [48].

**Fig 4.**
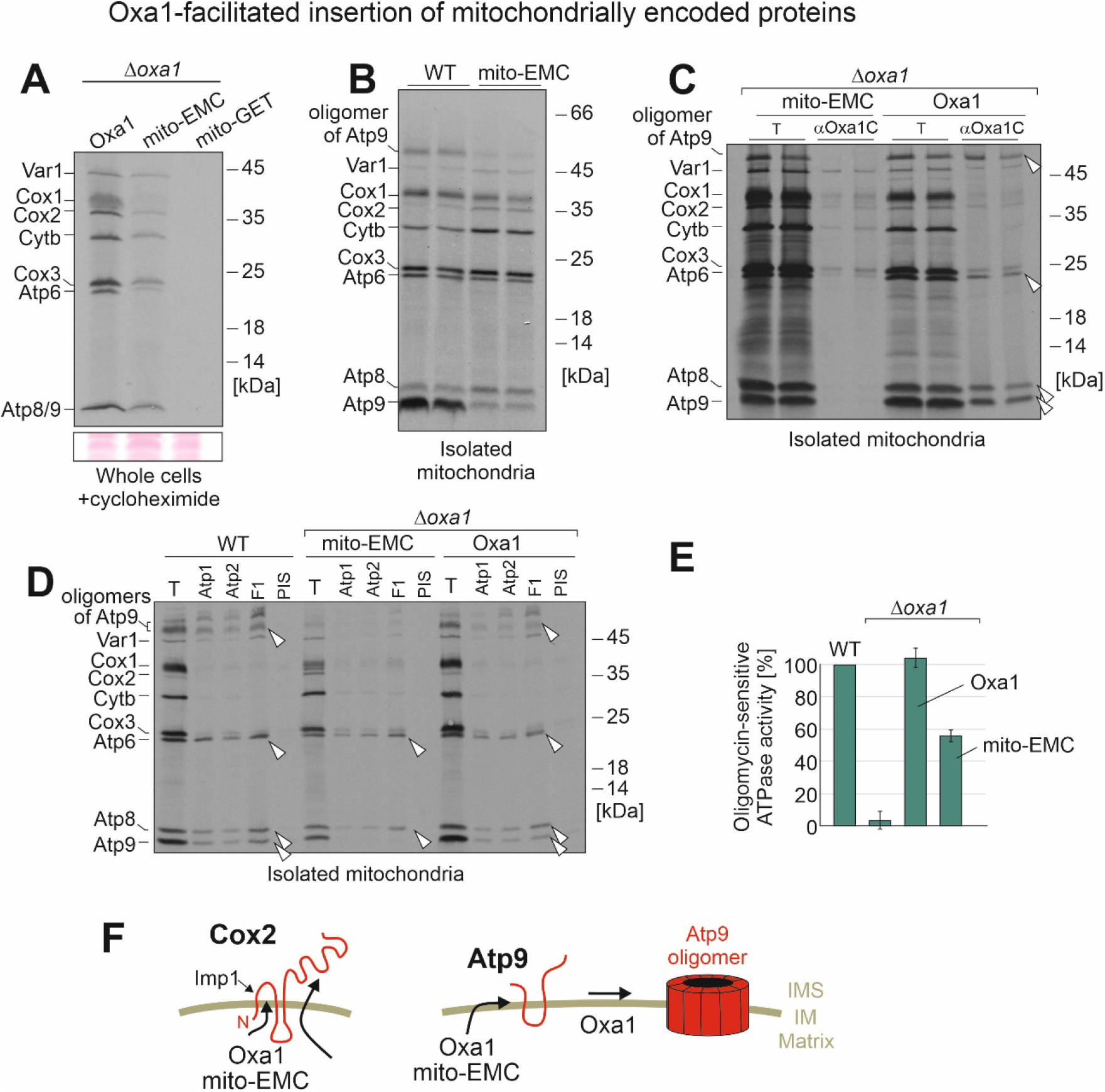
While mito-EMC mediates the insertion of Cox2, it fails to take over the Oxa1-facilitated assembly of the Atp9 oligomer. (**A**) *Δoxa1* mutants expressing Oxa1, mito-GET or mito-EMC were grown in galactose medium to log phase. Cytosolic translation was inhibited by cycloheximide. ^35^S-methionine was added to radiolabel mitochondrial protein synthesis. Cells were harvested and lysed. Proteins were precipitated by trichloroacetic acid and visualized by SDS-PAGE and autoradiography. (**B**) Mitochondria were isolated from the indicated strains and incubated in the presence of ^35^S-methionine in translation buffer for 30 min at 30°C. Mitochondria were washed and newly synthesized proteins visualized by autoradiography. (**C, D**) Mitochondrial translation products were radiolabeled for 30 min before mitochondria were lysed with Triton X-100. The extract was either directly loaded (T, equivalent to 10% total) or after immunoprecipitation with antibodies raised against the C-terminus of Oxa1 (αOxa1C), subunits of the ATPase or with preimmune serum (PIS). Arrowheads depict immunoprecipitated ATPase subunits. (**E**) The levels of oligomycin-sensitive ATPase were measured in isolated mitochondria and quantified. Shown are mean values of three replicates. (**F**) While mito-EMC is able to mediate the membrane insertion of Cox2, it does not efficiently promote the assembly of Atp9 into its oligomeric form. Thus, mito-EMC seems only to be able to take over some of the Oxa1-mediated functions in mitochondria.

The synthesis of mitochondrial translation products can be monitored in isolated mitochondria to which radiolabeled ^35^S-methionine is added [49]. Subunit 2 of cytochrome oxidase (Cox2) is initially produced as a precursor protein with an N-terminal leader sequence which is cleaved after membrane insertion. The accumulation of the Cox2 precursor is characteristic for *Δoxa1* mitochondria, owing to their membrane insertion deficiency [50]. Mitochondria containing mito-EMC instead of Oxa1 showed no or only low levels of Cox2 precursors (Fig. 4B). However, they showed strongly reduced amounts of Atp9, both in its monomeric form and in its SDS-resistant Atp9 oligomer. When mitochondria were lysed after the labeling reaction and Oxa1 and mito-EMC were isolated by immunoprecipitation with antibodies against the C-terminus of Oxa1, subunits of the ATPase were only co-isolated with Oxa1 but not with mito-EMC (Fig. 4C). When the labeling was stopped and isolated mitochondria were further incubated, monomeric Atp9 became converted into the oligomeric form in the presence of Oxa1, but to a much lesser degree in the mito-EMC mutant (Fig. S3). Moreover, immunoprecipitation of ATPase complexes with antibodies against Atp1, Atp2 or the F1-part of the ATPase copurified Atp6, Atp8 and Atp9 in Oxa1-containing mitochondria, but not in those of the mito-EMC mutant (Fig. 4D). From this we conclude that mito-EMC can insert Cox2 into the inner membrane. However, it does not efficiently promote the insertion and/or assembly of the Atp9 oligomer in the inner membrane. This is also confirmed by considerably diminished ATPase activity levels in mito-EMC mitochondria (Fig. 4E). Thus, whereas mito-EMC shares the insertase activity with Oxa1, it might be limited in its ability to interact with mitochondrial ATPase assembly factors (Fig. 4F).

## Discussion

The EMC complex of the ER was identified only rather recently [29]. The range of its physiological activities is not entirely clear, in part because its ability to insert proteins into the ER membrane overlaps with that of the Sec61 translocase [33]. The essential nature of the Sec61 complex makes it difficult to elucidate the EMC activity in the absence of that of the dominating translocon. The recently solved structure and the identification of a rather small Emc6/Emc3 core region of the EMC complex inspired us to try an *in vivo* reconstitution approach in the mitochondrial inner membrane. Our observations on this mito-EMC protein allow a number of conclusions:

1. Emc3 and Emc6 together indeed form a minimal insertase unit that is able to promote protein insertion into a lipid bilayer. Of course, in the native EMC complex, other subunits might add further activities and properties that are not reflected in the mito-EMC protein. However, the Emc6/Emc3 core region with their six transmembrane domains are sufficient to promote protein insertion. This observation confirms the insertion mechanism that was proposed on the basis of recent cryo-electron microscopic studies which revealed that a membrane-embedded hydrophobic groove formed by Emc3 and Emc6 constitutes a transient binding site that incorporates transmembrane stretches into a disturbed and locally thinned lipid bilayer [35-37, 51, 52].
2. Complexome profiling [30] and proteomic analysis of EMC-deficient cells [53] revealed that EMC substrates are transmembrane proteins, many of which span the ER membrane multiple times. Studies on reconstituted liposomes showed that while the EMC complex is able to integrate proteins of rather simple topology on its own [32], the assistance of the Sec61 translocon is often necessary for accurate topogenesis of multispanning proteins [31]. We observed that mito-EMC is able to mediate the insertion of the mitochondrial translation products which range from proteins of rather simple topology (Atp8 and Cox2) to multi-pass proteins with several transmembrane domains (Cox1, Cox3, cytochrome *b* and Atp6). Thus, our *in vivo* reconstitution indicates that, in principle, the EMC core is able to mediate the insertion of a broad range of membrane proteins in the absence of a Sec61-mediated insertion activity.
3. Oxa1 and YidC mediate the translocation of negatively charged protein segments, but largely fail to export positively charged regions [27, 46, 47]. The mito-EMC-mediated insertion adhered to the same properties, again supporting the conclusion that both insertases promote the same type of biochemical reaction.
4. In addition to its role as an insertase for membrane proteins, Oxa1 was proposed to facilitate the assembly of oligomeric complexes [54, 55]. Interestingly, we observed that the mito-EMC failed to promote efficient biogenesis of the Atp9 ring. Assembly of the F_0_ sector of the ATPase is a complicated process relying on a number of different assembly factors [56]. It was proposed that some of these are recruited by Oxa1 [38], consistent with our observations.

While the functional complementation of *Δoxa1* mutants by mito-EMC alone cannot prove a common evolutionary origin of these insertases, this observation strongly supports this assumption. We suggest that our findings demonstrate that the fundamental mechanisms by which proteins are integrated into the inner membrane of mitochondria and the ER are universal and the insertion process of membrane proteins adheres to the same principles despite the about three billion years of evolution since archaeal and bacterial lineages separated. This similarity may now be used to uncover fundamental principles of EMC function specifically and protein insertion more generally.

## Methods

### Yeast strains and plasmids

The yeast strains used in this study are either based YPH499 (MATa *ura3 lys2 ade2 trp1 his3 leu2*), or for the *Δcox18* experiment based on W303 (MATa *ura3 ade2 trp1 his3 leu2*). The *Δoxa1* [57] and *Δcox18* [12] strains used for this study were described before.

To generate the mito-GET and mito-EMC expression plasmids, the coding regions for the following protein stretches were amplified from genomic DNA and ligated into a pYX232 empty vector plasmid downstream of the *TPI* promoter by using the restriction sites *Xma*I and *Sal*I or *Sal*I and *Nco*I using a Gibson assembly protocol [58]: Oxa1(1-50), Get2(139-285), Oxa1(219-244), Get1(104-200), Oxa1(321-403) for mito-GET and Oxa1(1-50), Emc6(19-108), Oxa1(219-244), Emc3(1-197), Oxa1(321-403) for mito-EMC. For control, a pYX232 plasmid was generated that contained the entire Oxa1 reading frame (1-403). All plasmids were verified by sequencing.

Strains were grown at 30°C in minimal synthetic medium containing 0.67% yeast nitrogen base and 2% glucose, galactose or glycerol as carbon source. For plates, 1.5% of agar was added to media.

### Drop dilution assay

To test growth on plates, drop dilution assays were conducted. Yeast cells were grown in synthetic galactose media to mid log phase. After harvesting 2 OD (600 nm) of cells and washing with sterile water, a 1:10 serial dilution was prepared in sterile water. Equal amounts of the dilutions were dropped on agar plates to determine growth differences. Pictures of the plates were taken after 2 to 5 days of incubation.

### Growth curve assay

To test growth in liquid media, growth curves were performed. Yeast cells were grown in synthetic galactose media to mid log phase. After harvesting 2 OD (600 nm) of cells and washing with sterile water, cells of OD 0.1 were resuspended in the experimental media and transferred into a clear 96 well plate. Automated OD measurements were performed at 600 nm in the ELx808™ Absorbance Microplate Reader (BioTek®). ODs were measured for 72 h every 10 min at 30°C in technical triplicates.

### Isolation of mitochondria

To isolate crude mitochondria, yeast strains were cultivated in synthetic galactose media to mid log phase and harvested by centrifugation (5 min, 3000 g). After washing the pellets with water and centrifugation (5 min, 3000 g), the weight of the cell pellet was determined. Pellets were resuspended in 2 ml per g wet weight in MP1 (100 mM Tris, 10 mM DTT), incubated for 10 min at 30°C and centrifuged again (5 min, 3000 g). Pellets were washed with 1.2 M sorbitol and centrifuged (5 min, 3000 g) before resuspending in 6.7 ml per g wet weight in MP2 (20 mM KPi buffer pH 7.4, 1.2 M sorbitol, 3 mg per g wet weight zymolyase from Seikagaku Biobusiness) and incubated at 30°C for 60 min. The following steps were conducted on ice. After centrifugation (5 min, 2800 g), pellets were resuspended in 13.4 ml/g wet weight in homogenization buffer (10 mM Tris pH 7.4, 1 mM EDTA, 0.2% BSA, 1 mM PMSF, 0.6 M sorbitol) and a cooled glass potter was used to homogenize the sample with 10 strokes. After homogenization, the extract was centrifuged three times (5 min, 2800 g) while always keeping the mitochondria-containing supernatant. To pellet mitochondria, samples were centrifuged for 12 min at 17,500 g. Pellets were resuspended in SH buffer (0.6 M sorbitol, 20 mM HEPES pH 7.4). The concentration of the purified mitochondria was adjusted to 10 mg/ml protein. Aliquots were snap-frozen in liquid nitrogen and stored at -80°C.

### Import of radiolabelled precursor proteins into mitochondria

To prepare radiolabelled (^35^S-methionine) proteins for import experiments, the TNT® Quick Coupled Transcription/Translation Kit from Promega was used according to the instructions of the manufacturer. To determine the ability of proteins to be imported into mitochondria, *in vitro* import assays were conducted. Mitochondria were resuspended in a mixture of import buffer (500 mM sorbitol, 50 mM HEPES pH 7.4, 80 mM KCl, 10 mM Mg(OAc)_2_, 2 mM KPi) with 2 mM ATP and 2 mM NADH to energize them for 10 min at 25°C. The import reaction was started by addition of the radiolabelled lysate (1% final volume). Import was stopped after 30 min by transferring the mitochondria into cold SH buffer (0.6 M sorbitol, 20 mM HEPES pH 7.4) or 20 mM HEPES pH 7.4 (swelling conditions). The remaining precursors outside of the mitochondria were removed by protease treatment (PK) for 30 min. 2 mM PMSF was added to stop protein degradation. After centrifugation (15 min, 25000 g, 4°C), the supernatant was removed. Pellets were resuspended in SH buffer containing 150 mM KCl and 2 mM PMSF and centrifuged again (15 min, 25000 g, 4°C). Pellets were then lysed in sample buffer (2% sodium dodecyl sulfate, 10% glycerol, 50 mM dithiothreitol, 0.02% bromophenolblue, 60 mM Tris/HCl pH 6.8) and heated to 96°C for 5 min. Samples were run on a 16 % SDS-gel, blotted onto a nitrocellulose membrane and visualized with autoradiography.

### Labeling of mitochondrial translation products (*in organello* and *in vivo*)

Translation products were labeled in isolated mitochondria as described previously [59]. Mitochondria (100 µg protein) were incubated in translation buffer (0.6 M sorbitol, 150 mM KCl, 15 mM KH_2_PO_4_, 13 mM MgSO_4_, 0.15 mg/ml of all amino acids except methionine, 4 mM ATP, 0.5 mM GTP, 5 mM α-ketoglutarate, 5 mM phosphoenolpyruvate, 3 mg/ml fatty acid-free bovine serum albumin, 20 mM Tris/HCl pH 7.4) containing 0.6 U/ml pyruvate kinase and 10 µCi ^35^S-methionine. Samples were incubated for indicated timepoints at 30°C and labeling was stopped by addition of 25 mM unlabeled methionine. The samples were further incubated for 3 min to complete synthesis of nascent chains. Mitochondria were isolated by centrifugation, washed in 1 ml 0.6 M sorbitol, 20 mM HEPES/HCl, pH 7.4, lysed in 20 µl sample buffer and subjected to SDS-PAGE.

*In vivo* labeling of mitochondrial translation products was performed in whole cells in the presence of cycloheximide essentially as described [60]. Proteins were precipitated in the presence of 10% trichloroacetic acid, and precipitates washed with ice-cold acetone.

### ATPase activity assay

To determine the activity of the ATPase, 10 µl isolated mitochondria (100 µg protein) were dissolved in 90 µl lysis buffer (0.1% TX100, 300 mM NaCl, 5 mM EDTA, 10 mM HEPES/KOH pH 7.4) for 10 min and centrifuged (10 min, 16.000 rpm, 4°C). The supernatant was split into three aliquots. One remained untreated, one was treated with 70 µg/ml oligomycin to inhibit the F_o_/F_1_-ATPase function, and one was boiled at 96°C for 2 min to assess the phosphate background. 10 µl protein of each of these aliquots was transferred into 180 µl sample buffer (30 mM HEPES/KOH pH 7.4, 50 mM KCl, 5 mM MgSO_4_) before the reaction was started by addition of 10 µl 100 mM fresh ATP buffered in 300 mM HEPES/KOH pH 7.4. After 30 min at 25°C, 4 µl 70 µg/ml oligomycin and 330 µl of fresh solution 1 (160 mM ascorbic acid, 480 mM HCl, 3% SDS, 0.5% NH_4_MoO_4_) were added and samples were kept on ice for 10 min. Then, 500 µl solution 2 (70 mM sodium citrate, 2% sodium arsenite, 2% acetic acid) was added and samples were incubated at 25°C for 20 min. The absorbance was measured at 705 nm.

### Sequence analysis, alignments and phylogeny

Oxa1, Emc3, Get1 and Cox18 proteins from *Homo sapiens* and *Saccharomyces cerevisiae*, Alb3 and Alb4 from *Arabidopsis thaliana*, YidC from *Escherichia coli*, SpoIIIJ from Bacillus subtilis and *Methanocaldococcus jannaschii* DUF106 were used as seed sequences (Supplementary Data 1) to identify homologs using Diamond BLAST [61] against a database of 150 eukaryotes with complete genomes and RefSeq 2016 database [62]. A total of 460 unique homologs that had a minimum of 25% identity and an e-value of less than 1e-10 were chosen for further analysis (see Supplementary Data 2). In case of YidC and SpoIIIJ the first 50 hits were taken to reduce the number of homologs and compensate for the high number of prokaryotic representatives compared to eukaryotes. Maximum-likelihood trees were calculated using IQ-tree [63, 64] with standard parameters of a 1000 Ultrafast Bootstraps following an alignment performed using MAFFT [65] and trimming using TrimAl [66] and finally were rooted using MAD [67]. The final alignment is provided in Supplementary Data 3.

## Antibodies

Antibodies were raised in rabbits against proteins recombinantly expressed in bacteria and purified by affinity chromatography. The Oxa1-specific antibody was raised against the C-terminal region of Oxa1 that is present in both Oxa1 and mito-EMC. Antibodies against the mitochondrial ATPase were a gift from Marie-France Giraud from CNRS in Bordeaux.

## Data availability

All relevant data for this study is contained within the manuscript. Materials and strains are available from the authors.

## Acknowledgments

We thank Sabine Knaus, Andrea Trinkaus and Vanessa Scherer for technical assistance and Maya Schuldiner for discussions and comments on the manuscript. We are very grateful to Marie-France Giraud (CNRS, Bordeaux) for antibodies against ATPase subunits. This project was funded by grants from the Deutsche Forschungsgemeinschaft (DIP MitoBalance and HE2803/9-1) and the Landesschwerpunkt BioComp.

---

**Fig. S1.**
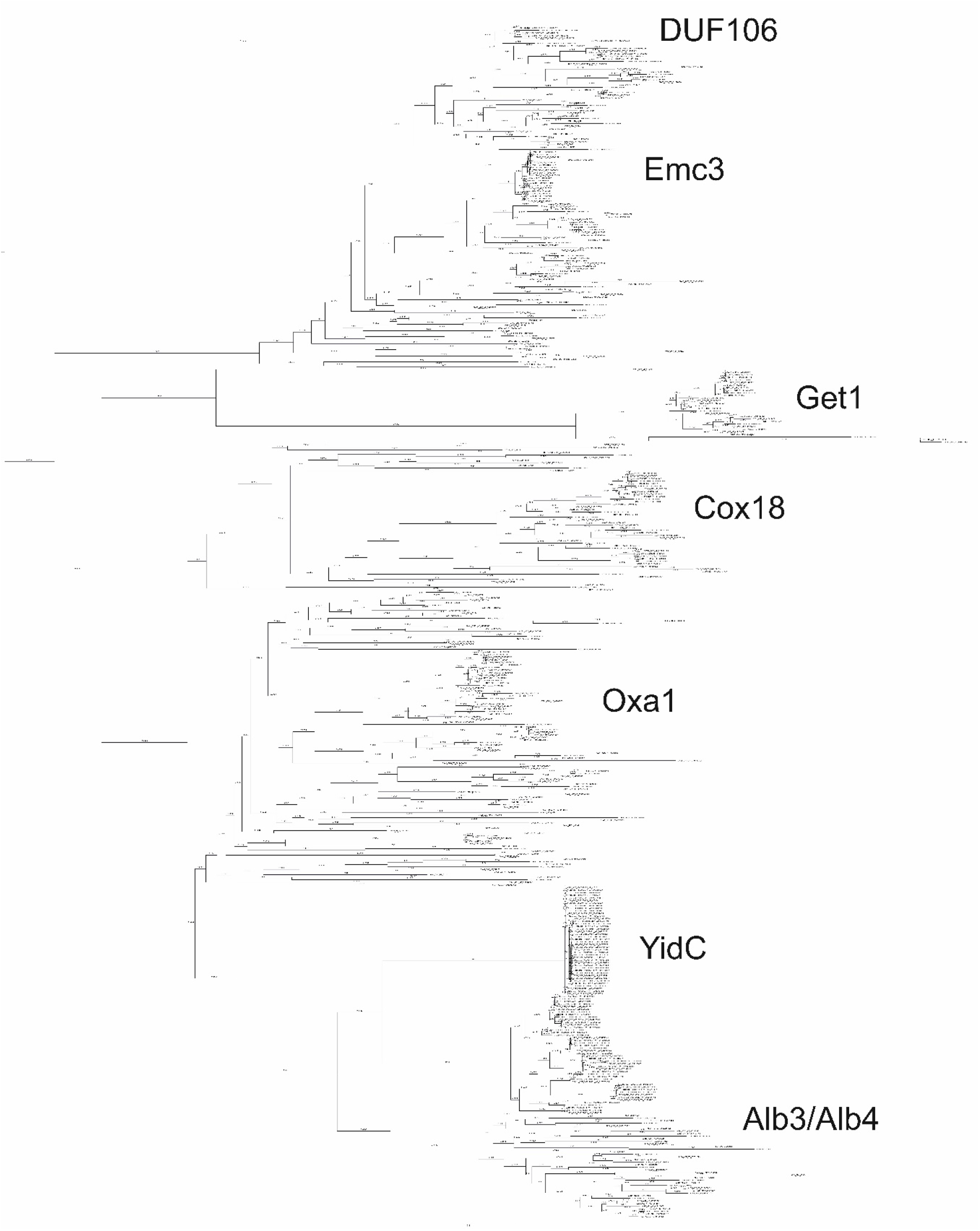
Complete rooted phylogenetic tree for homologs of DUF106, Emc3, Get1, Alb3/Alb4, Oxa1 and YidC. A rooted phylogenetic tree composed of sequence homologs of DUF106, Emc3, Get1, Alb3/Alb4, Oxa1 and YidC identified as described in the Methods section. The Boot strap values are shown at the nodes. The tree demonstrates a closer relationship between DUF106, Emc1 and Get1 compared to Cox18, YidC/Oxa1/Alb3. Alignments are provided in supplemental data set 3.

**Fig. S2.**
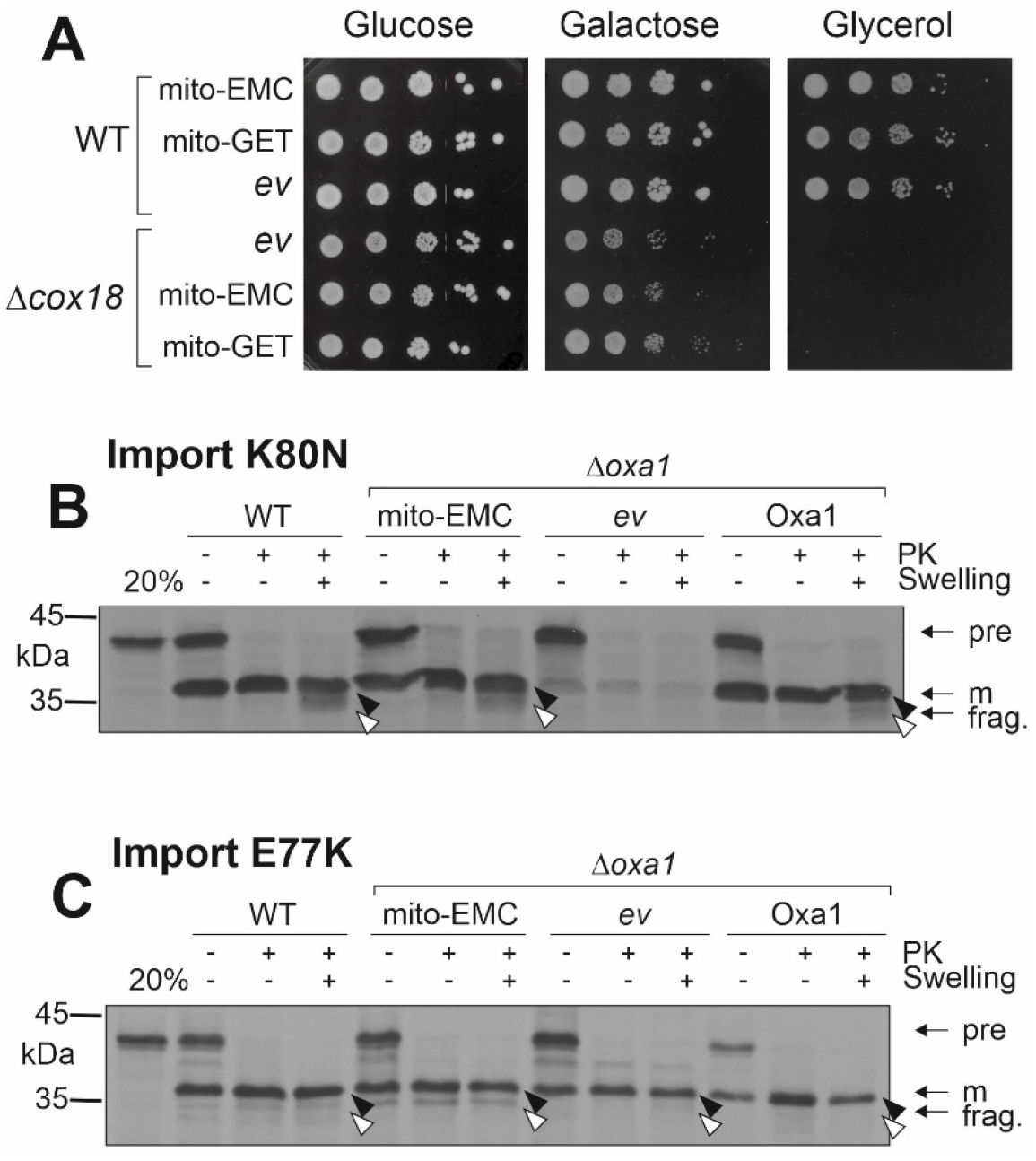
Neither mito-EMC nor mito-GET are able to take over the function of the Oxa1 homolog Cox18. (**A**) Wild type and *Δoxa1* cells were transformed with plasmids for the expression of mito-EMC or mito-GET or with an empty vector (*ev*) for control. Cells were grown on the indicated carbon sources. (**B, C**) Radiolabeled Su9-(1-112, K80N)-DHFR and Su9-(1-112, E77K)-DHFR were incubated with isolated mitochondria as described in Fig. 3C. For quantification, see Fig. 3D.

**Fig. S3.**
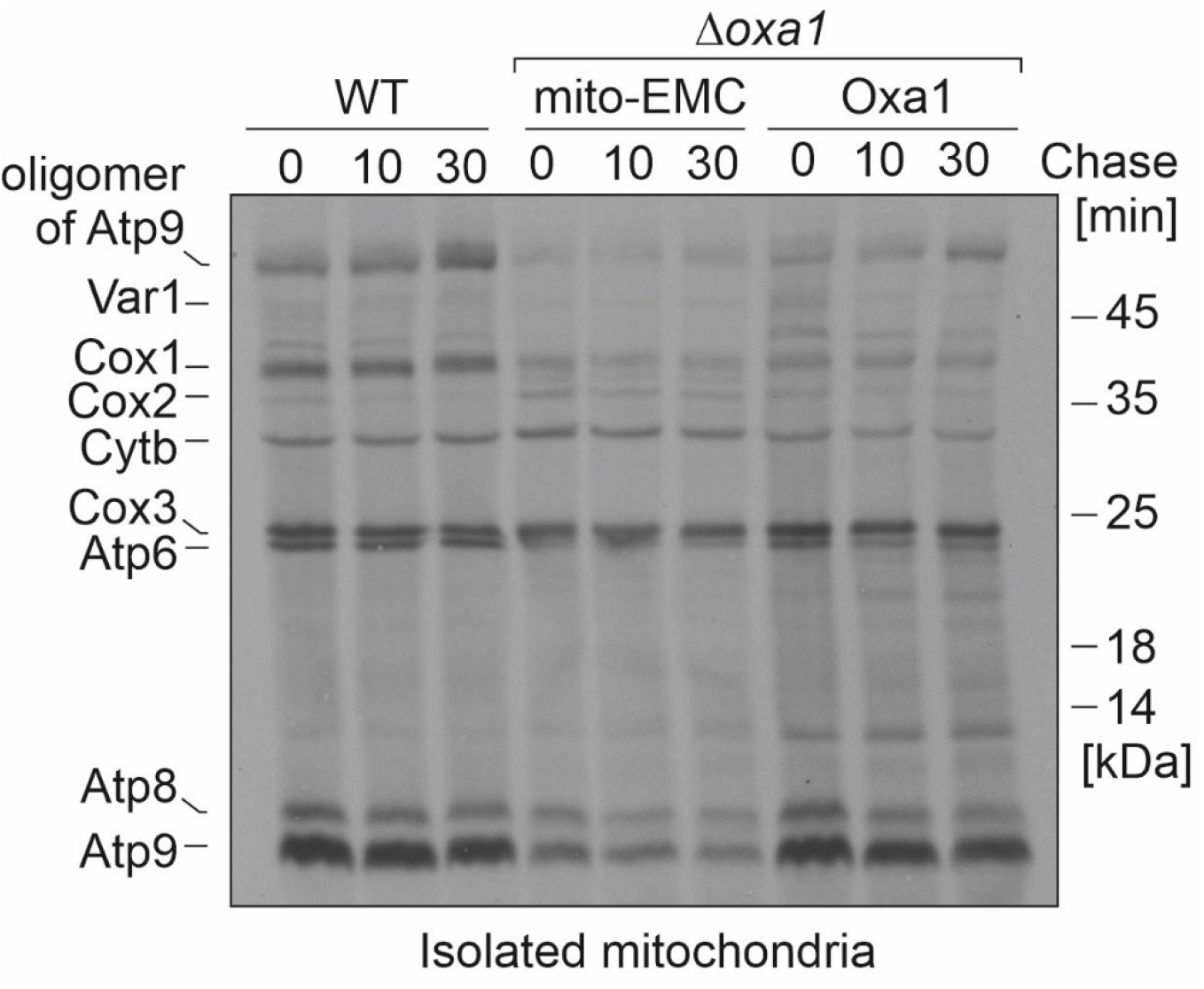
Assembly of ATPase is not efficiently promoted by mito-EMC. Newly synthesized proteins were radiolabeled in isolated mitochondria for 30 min. Radiolabeling was stopped by addition of an excess of cold methionine, and mitochondria were further incubated (chase) at 30°C for 0, 10 or 30 min.

